# TRACC-PHYSIO: Time-domain Resolution-Aligned Cross-Correlation to estimate PHYSIOlogical coupling and time delays in dynamic MRI

**DOI:** 10.1101/2025.08.01.668145

**Authors:** Adam M. Wright, Jianing Zhang, Yunjie Tong, Qiuting Wen

## Abstract

Physiological brain pulsations, primarily driven by cardiac and respiratory activity, play a key role in driving neurofluid circulation and waste clearance. Capturing the temporal dynamics of cardiac- and respiratory-driven brain pulsations (0.2-1.5 Hz) requires fast imaging with TRs near 100 ms, which is often unachievable in functional MRI or dynamic diffusion MRI. As a result, valuable physiological information remains hidden in these datasets. Here, we introduce TRACC-PHYSIO, a time-domain analytical framework designed to quantify physiological coupling and pulse time delays in dynamic MRI without requiring a fast acquisition. TRACC-PHYSIO uses cross-correlation to detect co-fluctuations between slowly sampled dynamic MRI data and simultaneously recorded physiological waveforms. It measures two key metrics: the peak Coupling Coefficient (peak CorrCoeff), quantifying the strength of co-fluctuations, and the TimeDelay, reflecting the relative arrival time of the physiological impulse in the brain with millisecond-level temporal resolution. The primary aim of this study is to validate TRACC-PHYSIO through systematic simulations that model realistic dynamic MR signals with mixed physiological components. We comprehensively evaluate TRACC-PHYSIO’s performance under a wide range of conditions, including varying cardiac-to-respiratory composition ratios, TRs, and acquisition times. Results demonstrate that TRACC-PHYSIO can robustly assess coupling strengths and time delays for both cardiac (TRACC-Cardiac) and respiratory (TRACC-Respiratory) components, even in datasets with long TRs up to 3 seconds. By enabling a reliable time-domain coupling analysis, TRACC-PHYSIO opens new avenues for revealing brain pulsation mechanisms and elucidating the physiological drivers of neurofluid dynamics in health and disease. This stimulation study provides a valuable reference for interpreting TRACC-PHYSIO results and understanding associated uncertainties in future applications.

**Highlights:**

- TRACC-PHYSIO is a time-domain method developed to estimate cardiac and respiratory coupling and their pulsation time delays in dynamic MRI without requiring high temporal resolution.
- TRACC-PHYSIO was validated through systematic simulations across varying physiological compositions and MR acquisition parameters.
- Results demonstrated that TRACC-PHYSIO reliably quantifies cardiac and respiratory components in dynamic MR signals.
- The stimulation study provides a useful reference for interpreting TRACC-PHYSIO results and understanding associated uncertainties in future applications.

## 1. Introduction

Physiological brain pulsations play a crucial role in driving neurofluid circulation and waste clearance (Iliff et al., 2013; Kiviniemi et al., 2016; Mestre et al., 2018; Nair et al., 2024; Wen et al., 2022; Wright et al., 2024). However, their underlying mechanisms and broad implications in aging and disease remain incompletely understood. Fast imaging with repetition times (TR) near 100 ms is typically required to resolve cardiac and respiratory temporal dynamics (Huotari et al., 2019; Kiviniemi et al., 2016; Raitamaa et al., 2021). These speeds are often unachievable in conventional dynamic MRI, such as functional MRI (fMRI) or dynamic diffusion-weighted imaging (dynDWI). As a result, valuable physiological information remains hidden in datasets with longer TRs. Here, we introduce a method named TRACC-PHYSIO, standing for Time-domain Resolution-Aligned Cross-Correlation, to estimate PHYSIOlogical coupling and time delays in dynamic MRI, which can estimate physiological coupling and pulse time delays in dynamic brain MRI without requiring fast imaging.

TRACC-PHYSIO detects time-domain co-fluctuations in undersampled brain data by leveraging information from simultaneous physiological recordings. Simply put, TRACC-PHYSIO performs a cross-correlation between a slow-sampled dynamic MRI signal and a fast-sampled physiological signal (cardiac or respiratory). The resulting cross-correlation waveform—termed TRACC-Cardiac or TRACC-Respiratory waveform—reveals the peak CorrCoeff and its corresponding TimeDelay, which represent the strength of coupling and the pulse arrival time in the brain relative to the physiological recording site (e.g., finger, chest). TRACC-PHYSIO provides a time-domain approach to study the cardiac and respiratory pulsations in slowly sampled data, where these pulsations are aliased in the frequency domain. It works because of the combination of natural variability in physiological rhythms and periodic MR sampling, which effectively samples different phases of the cardiac and respiratory cycles. This diverse phase coverage allows TRACC-PHYSIO to reliably estimate physiological coupling and timing, even in slow imaging acquisitions.

Recently, our group has applied the TRACC-PHYSIO approach to study the cardiac coupling and time delay in perivascular CSF using dynDWI (Wen et al., 2025, 2023), as well as pulse time delays between arteries and the superior sagittal sinus using fMRI (Wright et al., 2025). However, the performance of TRACC-PHYSIO hasn’t been thoroughly evaluated. Key questions remain: 1) How do the TR and acquisition time affect its performance? 2) How accurately can TRACC-PHYSIO assess coupling strength and time delays in the presence of noise and other physiological oscillations? To address these questions, first used a simplified simulation to confirm that TRACC-PHYSIO can accurately assess physiological coupling and time delays in a noise-free setting. We then performed in-depth simulations that reflect real-world physiological variability and noise conditions to systematically evaluate how varying TR, acquisition time, and compositions of cardiac and respiratory pulsations influence its accuracy. Finally, we interpret the physiological meaning of the two TRACC-PHYSIO output metrics—peak CorrCoeff and TimeDelay.

## 2. Methods

### 2.1. TRACC-PHYSIO analysis pipeline

TRACC-PHYSIO performs a cross-correlation between a dynamic MRI signal (typically sampled at TR>800 ms) and a simultaneously recorded physiological signal with a much higher sampling rate (i.e., 2.5 ms per sample, 400 Hz). It measures two key metrics: a Coupling Coefficient (CorrCoeff) that quantifies the strength of temporal coupling and a TimeDelay that reflects the relative arrival time of the physiological impulse in the brain compared to the physiological recording site (e.g., finger, chest).

In TRACC-PHYSIO, the MR signal is incrementally time-shifted in the smallest resolved step size based on the physiological sampling rate (2.5 ms for a 400 Hz signal), relative to the physiological signal. After each shift, a correlation is computed between the MR signal and the aligned physiological segment (Figure 1A). The resulting cross-correlation waveform—termed TRACC-Cardiac or TRACC-Respiratory waveform—reveals the peak CorrCoeff and its corresponding TimeDelay (Figure 1B). The stepwise cross-correlation and resulting TRACC-Cardiac waveform construction are summarized in Supplemental Video 1. Note that the final cross-correlation waveform (TRACC-Cardiac or TRACC-Respiratory) has a high temporal resolution equivalent to the physiological sampling rate (400 Hz).

**Figure 1:**
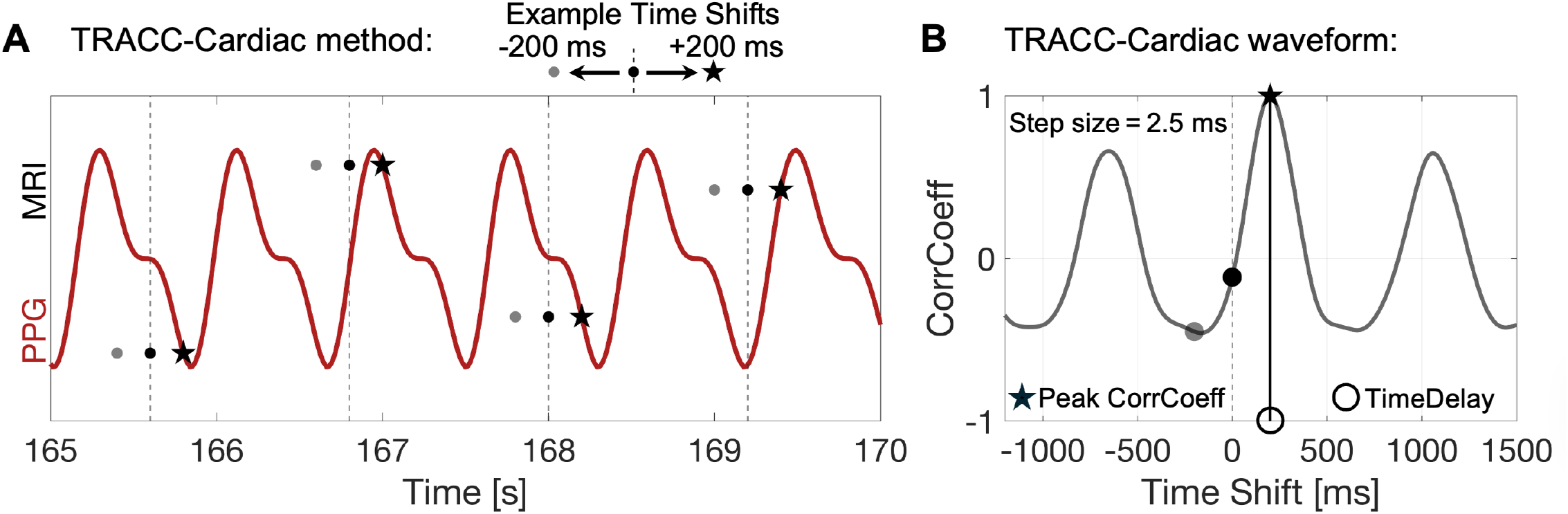
Overview of the TRACC-PHYSIO method to measure peak cardiac coupling and pulse time delay. **A**. The MRI timeseries (solid black dots) is shifted in increments of ±2.5 ms, and the correlation with the physiological signal (finger PPG) is computed at every time step. Examples of the time-shifted MR signal are illustrated at three shifts: -200 ms (gray point), 0 ms (black point), and +200 ms (black star). **B**. The resulting cross-correlation curve captures the degree of co-fluctuation between MR signal and PPG signal across time shifts. The three example shifts from panel A (gray point, black point, black star) are marked on the curve to illustrate their corresponding correlation values (CorrCoeff). The peak CorrCoeff and its associated time shift (TimeDelay) indicate when the strongest coupling occurs. In this example, a peak CorrCoeff of 1 indicates perfect coupling between the MR and PPG signals, with a TimeDelay of +200 ms, indicating that the MR signal leads the PPG signal by 200 ms. Note that the peak CorrCoeff is higher than the neighboring side peaks in the TRACC-Cardiac waveform; variability in heart rate leads to reduced correlation when aligning with pulses that are offset by one cycle. **Abbreviations:** PPG – photoplethysmography.

### 2.2. Simulation signal design

#### 2.2.1. Physiological signals

Synthetic finger photoplethysmography (PPG) signals were generated by combining a smooth sine wave with a high-frequency asymmetric component to best represent a cardiac beat with a sharp systolic upstroke, dicrotic notch, and gradual diastolic decay with a sampling frequency of 400 Hz. The heart rate (HR) and heart rate variability (HRV) could be adjusted to represent any combination of physiologically plausible parameters in future simulations. Supplemental Figure 1A illustrates a finger PPG waveform and frequency spectrum of a simulated PPG signal with an HR of 70 beats-per-minute and HRV characterized by a standard deviation of beat-to-beat intervals of 80 ms.

Similarly, synthetic chest respiratory belt signals were generated using a sinusoidal waveform with an asymmetry to reflect the natural variability in inspiratory and expiratory durations with a sampling frequency of 400 Hz. The respiratory rate (RR) and respiratory rate variability (RRV) could be adjusted to reflect any physiologically plausible breathing patterns. Supplemental Figure 1B illustrates a respiratory waveform and frequency spectrum of a simulated respiratory signal with an RR of 14 breaths per minute and RRV characterized by a standard deviation of breath-to-breath intervals of 300 ms.

#### 2.2.2. MR signals

Synthetic MR signals composed of cardiac and respiratory components were generated by combining modified physiological waveforms. The cardiac component of the MR signal was assigned the same polarity as the physiological cardiac signal, such that an increase in the cardiac waveform (i.e., finger PPG) corresponded to increases in the MR cardiac component. In contrast, the respiratory component was given the opposite polarity, such that increases in the respiratory waveform (i.e., chest belt) corresponded to decreases in the MR respiratory component. These polarity relationships match previous observations from dynDWI studies (Wen et al., 2025, 2023; Zhang et al., 2025). The cardiac and respiratory components were scaled to create MR signals with varying physiological compositions. All MR signals were initially generated at the physiological signal sampling rate of 400 Hz and an acquisition time of 600 s, then downsampled and truncated to simulate different TRs and acquisition times.

### 2.3. Simulation design

#### 2.3.1. Simplified proof of concept simulation

In a simplified simulation, a noise-free MR signal with varying TRs (TR = 0.05, 0.8, 1.6, and 2.4 s) and an acquisition time of 300 s was simulated to contain equal-amplitude cardiac and respiratory components (C:R=1:1). The MR cardiac component was time-shifted to lead the cardiac physiological signal by 200 ms. Then, TRACC-Cardiac was applied to estimate the peak CorrCoeff and TimeDelay between the two signals. This simulation provides a concise evaluation of how TRACC-Cardiac performs across different TRs.

#### 2.3.2. In-depth simulations with varying MR and physiological parameters

In-depth simulations were conducted to evaluate TRACC-PHYSIO’s performance under complex, real-world conditions, incorporating the following factors: **(1) MR signal noise**: Random noise was added to the MR signals to achieve a temporal signal-to-noise ratio of 20 dB. **(2) Cardiac-to-respiratory composition**: MR signals were simulated with three different cardiac-to-respiratory ratios (C:R), including 2:1, 1:1, and 1:2. **(3) TR and acquisition time**: Simulations spanned TR values from 0.05 to 3 s (in 0.025 s increments) and acquisition times from 60 to 580 s (in 20 s increments) to assess their impact on TRACC performance. **(4) Physiological variability**: Cardiac and respiratory waveforms were generated with varying HR, HRV, RR, and RRV, based on predefined ranges listed in Table 1. **(5) Variations in physiological time shift:** Randomized time shifts between MR and physiological signals were introduced to simulate natural biological variability in TimeDelay.

**Table 1:**
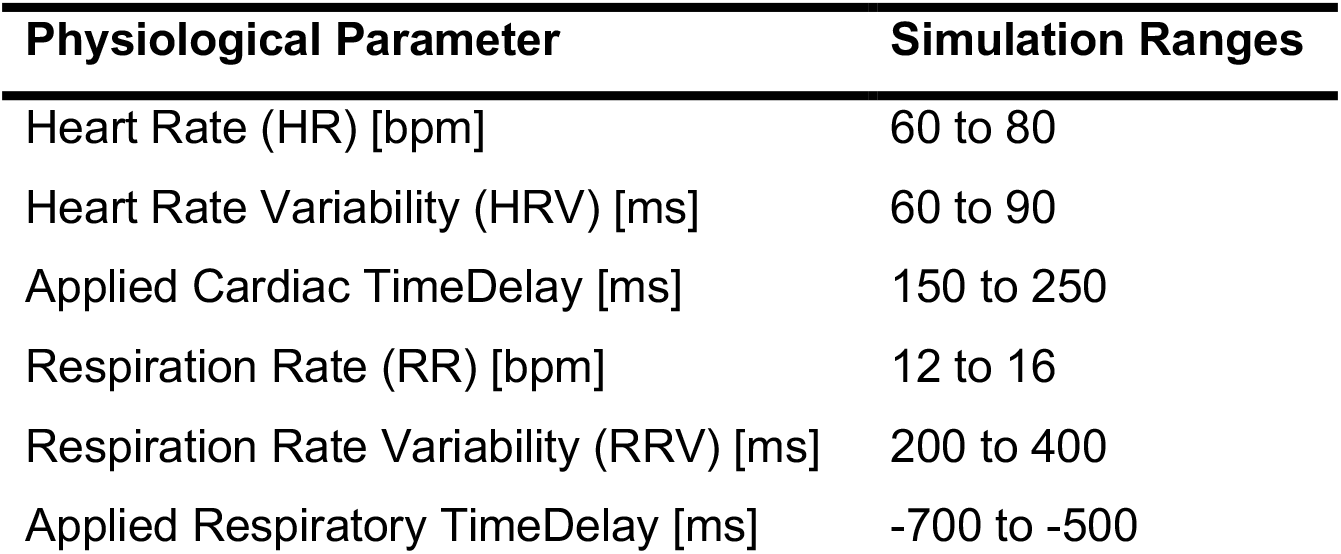
Physiological parameter ranges used in simulations. **Note:** Heart rate variability (HRV) and respiration rate variability (RRV) were defined as the standard deviation of beat-to-beat intervals and breath-to-breath intervals, respectively. **Abbreviations:** bpm – beats per minute or breaths per minute.

| Physiological Parameter | Simulation Ranges |
| --- | --- |
| Heart Rate (HR) [bpm] | 60 to 80 |
| Heart Rate Variability (HRV) [ms] | 60 to 90 |
| Applied Cardiac TimeDelay [ms] | 150 to 250 |
| Respiration Rate (RR) [bpm] | 12 to 16 |
| Respiration Rate Variability (RRV) [ms] | 200 to 400 |
| Applied Respiratory TimeDelay [ms] | -700 to -500 |

For each MR signal configuration (defined by a particular C:R ratio, TR, and acquisition time), 5,000 permutations were performed. In each permutation, randomized HR, HRV, RR, and RRV were used to generate the physiological signals to mimic an individual’s variability, the ground-truth TimeDelay was randomized, and random noise was added to the MR signal. TRACC-PHYSIO was then applied to estimate the peak CorrCoeff and TimeDelay between the MR signal and each physiological signal (TRACC-Cardiac and TRACC-Respiratory). TimeDelay error was computed as the difference between the ground-truth and the TRACC-estimated TimeDelay.

## 3. Results

### 3.1. TRACC-PHYSIO demonstrated high performance across variable TRs in simplified simulations

In the noise-free, simplified simulation with equal amplitude physiological components (C:R=1:1), TRACC-Cardiac produced consistent results across TRs ranging from 0.05 to 2.4 s (Figure 2). At TR=0.05s, it identified a peak CorrCoeff of 0.66 and a TimeDelay of 200 ms (Figure 2A). With longer TRs of 0.8, 1.6, and 2.4 s, TRACC-Cardiac yielded peak CorrCoeffs between 0.65-0.68 and TimeDelays between 200-202.5 ms (Figure 2B–D), differing no more than 0.02 in peak CorrCoeff and 2.5 ms in TimeDelay from the TR = 0.05 s baseline. This simulation demonstrates that TRACC-Cardiac’s accuracy and robustness even at long TRs.

**Figure 2:**
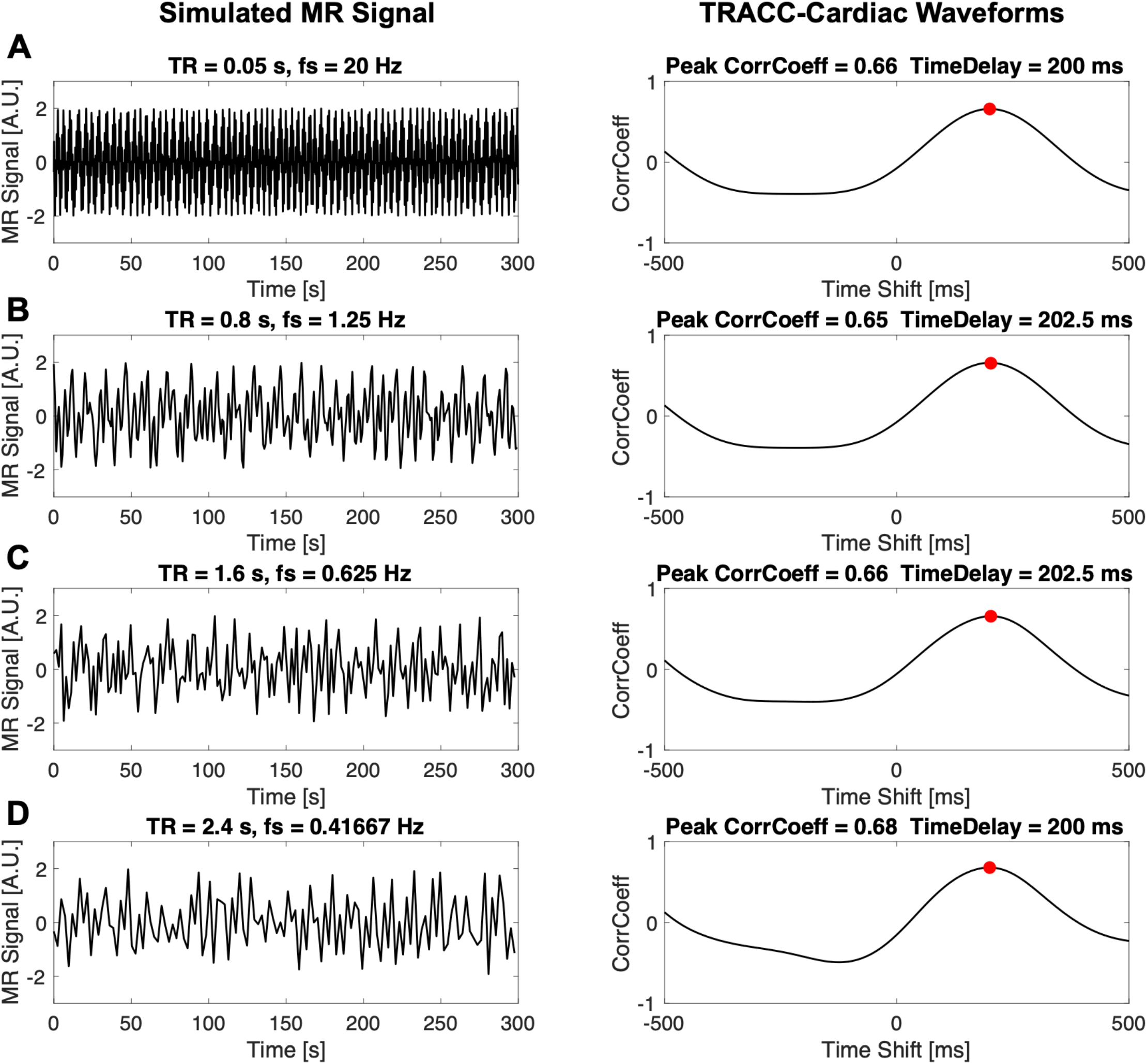
TRACC-PHYSIO demonstrated high performance across variable TRs in simplified simulations. TRACC-Cardiac was applied to simulated MR signals with equal cardiac and respiratory components with a ground-truth cardiac TimeDelay of 200 ms. Peak CorrCoeff and TimeDelay estimates were consistent across short (**A**: TR = 0.05 s) and longer TRs (**B**: TR = 0.8 s; **C**: TR = 1.6 s; **D**: TR = 2.4 s).

### 3.2. TRACC-PHYSIO demonstrated robustness in in-depth simulations

In simulations where the physiological coupling being solved for dominates the MR signal (i.e., C:R=2:1 for TRACC-Cardiac and C:R=1:2 for TRACC-Respiratory), TRACC-PHYSIO yielded high peak CorrCoeffs (>|0.8|) and accurately measured TimeDelays across various TRs and acquisition times (Figure 3). Across all simulations, both peak CorrCoeff and TimeDelay measures had no mean error bias.

**Figure 3:**
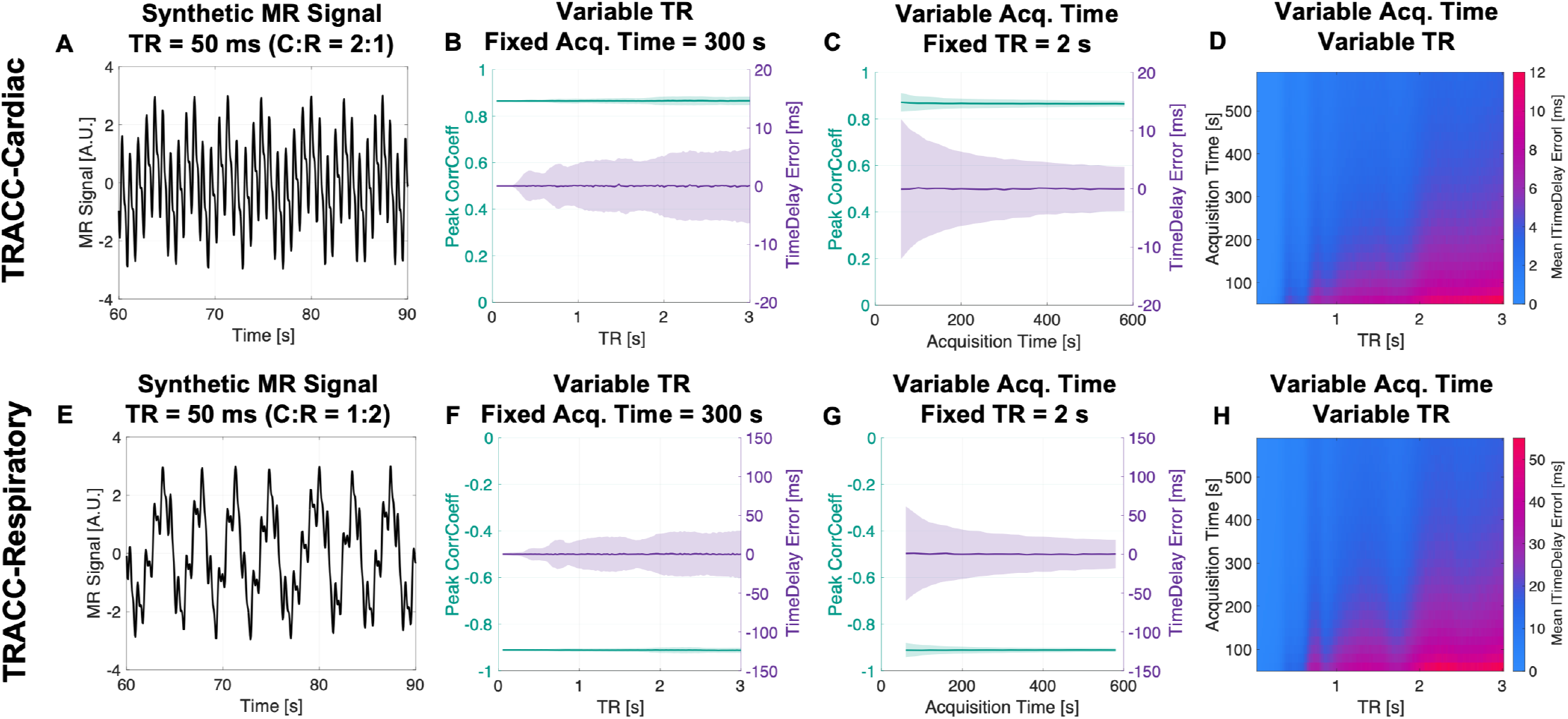
TRACC-PHYSIO demonstrated robust performance across varying TRs and acquisition times when the target physiological component is dominant in the MR signal. Top row (**A-D**): TRACC-Cardiac results for MR signals with C:R=2:1. Bottom row (**E-H**): TRACC-Respiratory results for MR signals with C:R=1:2. From left to right: (**A & E**) Example synthetic MR signals with TR = 50 ms. (**B & F**) The mean (solid line) and standard deviation (shaded) of the peak CorrCoeff and TimeDelay error with increasing TRs and a fixed acquisition time of 300 s. (**C & G**) The mean (solid line) and standard deviation (shaded) of the peak CorrCoeff and TimeDelay error with increasing acquisition time and a fixed TR of 2 s. (**D & H**) Heatmap of the mean absolute TimeDelay error across all combinations of TRs and acquisition times. **Note:** 5000 permutations were completed for each combination of TR and acquisition time. **Abbreviations:** C:R – cardiac-to-respiratory ratios.

In both TRACC-Cardiac and TRACC-Respiratory, across all TRs with a fixed acquisition time of 300 s, the mean peak CorrCoeff and mean TimeDelay error remained consistent, with values of 0.87 and 0 ms for TRACC-Cardiac and -0.91 and 0 ms for TRACC-Respiratory (Figure 3B, 3F). In TRACC-Cardiac, the standard deviation of the TimeDelay error increased starting at TR = 0.275 s, just before cardiac aliasing would occur in the frequency domain; however, the errors never exceeded ±6.5 ms. In TRACC-Respiratory, standard deviation increased at TR = 0.3 s and again at TR = 0.6 s, but the errors never exceeded ±30 ms. With a fixed TR = 2 s and increasing acquisition time, the mean peak CorrCoeff and mean TimeDelay error remained consistent in both TRACC-Cardiac and TRACC-Respiratory, and the measures’ standard deviation decreased as the acquisition time increased (Figure 3C, 3G). The mean absolute TimeDelay errors for TRACC-Cardiac and TRACC-Respiratory across all combinations of TR and acquisition time are summarized in Figures 3D and 3H, demonstrating that TimeDelay errors remained low across most acquisition parameters and were higher in simulation with longer TR and shorter scan durations.

In simulations where the physiological coupling being solved for was non-dominant in the MR signal (i.e., C:R=1:2 for TRACC-Cardiac and 2:1 for TRACC-Respiratory), TRACC-PHYSIO yielded moderate peak CorrCoeffs (∼|0.4 − 0.5|) and maintained accurate TimeDelay estimates across various TRs and acquisition times (Figure 4). As expected, uncertainty in both peak CorrCoeff and TimeDelay was higher compared to simulations with dominant coupling (Figure 4B vs Figure 3B, 4F vs 3F). Slight biases in TimeDelays emerged at short acquisition times (< 200 s) with a TR of 2 s (Figure 4C), but these biases disappeared with longer acquisition times. Heatmaps of mean absolute TimeDelay errors across TR and acquisition time combinations (Figures 4D and 4H) demonstrate that the most accurate performance was achieved with shorter TRs and longer acquisition times.

**Figure 4:**
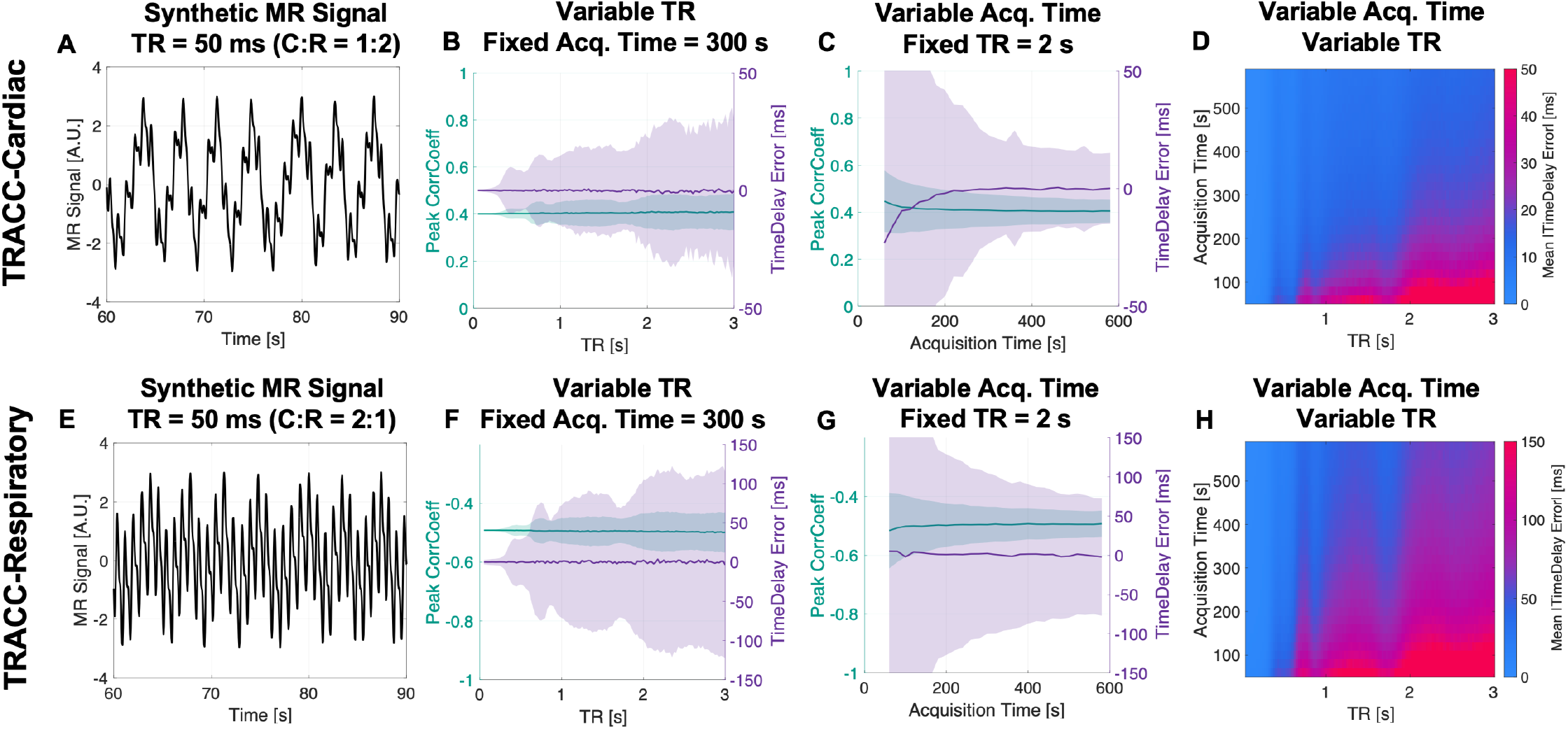
TRACC-PHYSIO maintained reasonable accuracy even when the target physiological component is non-dominant in the MR signal. Top row (**A-D**): TRACC-Cardiac results for MR signals with C:R=1:2. Bottom row (**E-H**): TRACC-Respiratory results for MR signals with C:R=2:1. From left to right: (**A & E**) Example synthetic MR signals with TR = 50 ms. (**B & F**) The mean (solid line) and standard deviation (shaded) of the peak CorrCoeff and TimeDelay error with increasing TRs and a fixed acquisition time of 300 s. (**C & G**) The mean (solid line) and standard deviation (shaded) of the peak CorrCoeff and TimeDelay error with increasing acquisition time and a fixed TR of 2 s. (**D & H**) Heatmap of the mean absolute TimeDelay error across all combinations of TRs and acquisition times. **Note:** 5000 permutations were completed for each combination of TR and acquisition time. **Abbreviations:** C:R – cardiac-to-respiratory ratios.

In simulations where the physiological components had equal amplitude in the MR signal (i.e., C:R=1:1), TRACC-PHYSIO yielded peak CorrCoeffs in the range of |0.6 − 0.8| and accurately measured TimeDelays across various TRs and acquisition times, which are fully summarized in Supplemental Figure 2. Across all simulations, the errors observed in TRACC-PHYSIO were between those observed in solving for the dominant and non-dominant physiological components.

### 3.3. TRACC-PHYSIO’s performance in five-minute scans

The simulation results in a typical acquisition time of 300 s at representative TRs and varying physiological signal components are summarized in Table 2. In both TRACC-Cardiac and TRACC-Respiratory, absolute peak CorrCoeff values progressively decreased as the relative contribution of the target physiological signal decreased. Using TRACC-Cardiac in cardiac-dominant signals with a long TR of 3 s, the mean absolute TimeDelay error was 5.1 ms. Errors were slightly higher when using TRACC-Respiratory in respiratory-dominant signals with a long TR of 3 s, yielding a mean absolute TimeDelay error of 24.0 ms. When solving for non-dominant physiological components, the results yielded higher TimeDelay errors: 20.4 ms for TRACC-Cardiac and 92.8 ms for TRACC-Respiratory. In all scenarios, TimeDelay errors progressively decreased with shorter TRs.

**Table 2:** Summary of TRACC-PHYSIO performance in five-minute scans. Each row summarizes simulation results of 5000 permutations for a specific cardiac-to-respiratory (C:R) physiological composition and TR. For each permutation, the applied TimeDelay was randomly selected between 150 and 250 ms (mean = 200 ms) for cardiac and between -700 and -500 ms for respiration (mean = -600 ms), with the heart rate, heart rate variability, respiratory rate, and respiratory rate variability randomized within a physiologically plausible range. Gaussian noise was added to achieve a temporal signal-to-noise ratio of 20 dB.

| Cardiac-to-Respiratory Ratio (C:R) | TR [s] | TRACC-Cardiac |  | TRACC-Respiratory |  |
| --- | --- | --- | --- | --- | --- |
|  |  | Peak CorrCoeff Mean (5-95%) | Absolute TimeDelay Error (ms) Mean (5-95%) | Peak CorrCoeff Mean (5-95%) | Absolute TimeDelay Error (ms) Mean (5-95%) |
| 2:1 | 0.05 | 0.87 (0.86, 0.87) | 0.0 (0, 0) | -0.49 (-0.49, -0.49) | 2.2 (0, 5.0) |
|  | 0.8 | 0.87 (0.85, 0.88) | 2.8 (0, 7.5) | -0.49 (-0.56, -0.43) | 51.5 (2.5, 140.0) |
|  | 1 | 0.87 (0.85, 0.88) | 2.9 (0, 7.5) | -0.49 (-0.56, -0.42) | 54.6 (5.0, 145.0) |
|  | 2 | 0.87 (0.84, 0.89) | 4.0 (0, 10.0) | -0.50 (-0.59, -0.39) | 78.1 (5.0, 210.0) |
|  | 3 | 0.87 (0.84, 0.89) | 5.1 (0, 12.5) | -0.50 (-0.61, -0.38) | 92.8 (7.5, 232.5) |
| 1:1 | 0.05 | 0.66 (0.66, 0.66) | 0.0 (0, 0) | -0.75 (-0.75, -0.75) | 1.1 (0, 2.5) |
|  | 0.8 | 0.66 (0.61, 0.70) | 5.5 (0, 15.0) | -0.75 (-0.78, -0.71) | 25.2 (2.5, 70.0) |
|  | 1 | 0.66 (0.61, 0.70) | 5.8 (0, 15.0) | -0.75 (-0.78, -0.71) | 26.9 (2.5, 75.0) |
|  | 2 | 0.66 (0.58, 0.73) | 8.3 (0, 20.0) | -0.75 (-0.80, -0.69) | 39.2 (2.5, 102.5) |
|  | 3 | 0.66 (0.58, 0.73) | 9.7 (0, 25.0) | -0.75 (-0.80, -0.69) | 46.7 (2.5, 117.5) |
| 1:2 | 0.05 | 0.40 (0.40, 0.40) | 0.0 (0, 0) | -0.91 (-0.91, -0.91) | 0.7 (0, 2.5) |
|  | 0.8 | 0.40 (0.33, 0.48) | 11.1 (0, 30.0) | -0.91 (-0.92, -0.90) | 13.2 (0, 35.0) |
|  | 1 | 0.40 (0.32, 0.48) | 11.6 (0, 30.0) | -0.91 (-0.92, -0.90) | 13.8 (0, 37.5) |
|  | 2 | 0.41 (0.29, 0.52) | 16.5 (2.5, 42.5) | -0.91 (-0.93, -0.89) | 20.2 (2.5, 52.5) |
|  | 3 | 0.41 (0.27, 0.53) | 20.4 (2.5, 52.5) | -0.91 (-0.93, -0.89) | 24.0 (2.5, 60.0) |

## 4. Discussion

Through simulations, we demonstrated that TRACC-PHYSIO can effectively assess physiological coupling and pulsation arrival timing in dynamic MR. We comprehensively characterized the performance of TRACC-PHYSIO in MR signals with mixed physiological components. Below, we interpret the peak CorrCoeff and TimeDelay metrics and discuss how MR acquisition parameters and the physiological composition influence TRACC-PHYSIO’s performance.

The peak CorrCoeff quantifies the degree to which the MRI signal dynamics are coupled with physiological fluctuations. Its square (CorrCoeff^2^ or r^2^) reflects the proportion of MR signal variance explained by the physiological input. A higher CorrCoeff indicates stronger co-fluctuation and provides a valuable metric for assessing physiological contributions to MR signals, particularly in long TR acquisitions. In neurofluid imaging modalities such as fMRI and dynDWI, long TRs (>800 ms) lead to aliasing of cardiac and respiratory frequencies, hindering their spectral assessment. Consequently, the physiological drivers across different brain regions are still under debate (Bohr et al., 2022). Our simulation results demonstrate that TRACC-Cardiac and TRACC-Respiratory can be applied to these MR signals, with peak CorrCoeff offering a robust measure of cardiac and respiratory contributions. Recent application of TRACC-PHSYIO suggests that parenchymal fluid dynamics are strongly coupled to respiration, whereas the fluid motion within the subarachnoid space is more closely linked to cardiac pulsatility (Zhang et al., 2025).

The TimeDelay reflects the latency between the physiological pulsation arrival in the brain and the peripheral physiological signal. A key advantage of TRACC-derived TimeDelay is its high millisecond-level temporal resolution, which matches the sampling rate of the physiological signal recordings. This fine resolution enables the detection and quantification of fast-traveling physiological waves to and within the brain. For example, TRACC-PHYSIO has revealed a 244 ms delay in cardiac pulsation between brain and finger (Wen et al., 2023), an 84 ms cardiac transmission from the subarachnoid space to the cortical surface (Wen et al., 2025), and a ∼80 ms delay from the cerebral arteries of the Circle of Willis to the superior sagittal sinus (Wright et al., 2025). The high temporal precision also allows for assessment of age-related changes in pulse propagation. Another advantage of the TRACC-derived TimeDelay is that it is measured in the time domain, preserving the true timing relationships between signals. In contrast, phase delays measured with retrospective rebinning approaches can introduce phase distortions by forcing cycles of variable length (from natural physiological variation) into a fixed phase scale, which may cause temporal offset errors. However, TRACC-PHYSIO requires a different consideration: to avoid misalignment with neighboring pulses under noisy conditions, the TimeDelay search window should be restricted to one-half of a physiological period and empirically centered on a physiologically plausible time based on group averages. Given the advantages and specific considerations of TRACC-PHYSIO, it holds great promise for broad applications to advance our understanding of neurofluid pulse timing and its alterations in aging and disease.

Our simulations indicate that both TR and acquisition time affect the accuracy of peak CorrCoeff and TimeDelay measures. As expected, short TR and longer acquisition time improve accuracy. Overall, peak CorrCoeff was very stable across all MR parameters (Supplemental Figure 3 and 4). TimeDelays errors in both TRACC-Cardiac and TRACC-Respiratory increased around a TR of 300 ms, coinciding with the onset of cardiac aliasing. Under typical MR acquisition paradigms (TRs of 1-3 seconds; 5-minute acquisition), TimeDelays remained reasonably accurate. Specifically, when TRACC-Cardiac was applied to a C:R=2:1 signal, the mean absolute TimeDelay errors were 2.9 and 4.0 ms at TRs of 1 and 2 seconds, respectively. For TRACC-Respiratory, with a C:R=1:2 signal, the corresponding errors were 13.8 and 20.2 ms. The TimeDelay errors were independent of the ground-truth TimeDelay (results not shown). For representative cardiac and respiratory TimeDelays of 200 ms and -600 ms, relative errors ranged from 1.5-2% for TRACC-Cardiac and 2.3-3.4% for TRACC-Respiratory, across TRs of 1-2 s. The higher error in TRACC-Respiratory likely stems from the lower respiratory frequency: over the same scan duration, respiratory cycles are 4-6 fold fewer than cardiac cycles, yielding less precise signal alignment.

Additionally, the physiological composition influences the accuracy of TRACC-PHYSIO results, particularly in TimeDelay estimates. When solving for the non-dominant physiological components, relative TimeDelay errors increased when compared to solving for the dominant physiological component. This suggests that peak CorrCoeff, which reflects the level of physiological dominance, can serve as an indicator of confidence in the TimeDelay measurement. Supporting this, our supplementary regression analysis showed a significant association between TimeDelay error and peak CorrCoeff (see Supplemental Tables 1 & 2). In summary, CorrCoeff not only reflects physiological coupling strength (see Supplemental Figures 5 & 6), but also provides a measure of certainty of the TimeDelay measurement. Importantly, despite greater variability in TimeDelay for non-dominant components, TRACC-PHYSIO introduced no systematic bias in either peak CorrCoeff or TimeDelay estimates. The symmetric error distribution ensures valid and interpretable group-level results, with estimates converging towards the true values.

The TRACC-PHYSIO method contains certain limitations. First, it relies on high-quality peripheral physiological recordings, which can be challenging to acquire consistently. Poor physiological signals reduce the TRACC-PHYSIO accuracy, making rigorous quality control of the physiological data essential. Second, the peak CorrCoeff is influenced by the waveform shape similarity between peripheral physiological signals and MR signals. Variations in MR waveform shape may arise from system-specific transfer functions in response to physiological impulses. While these transfer functions are not well characterized for many neurofluid compartments, a better understanding of them represents an opportunity for future refinement of the TRACC-PHYSIO framework. Third, the TRACC-PHYSIO method involves aligning MR slice timing with the reference physiological signal; this alignment restricts the application of motion correction, as it would disrupt slice timings and this alignment. In the future, it may be possible to apply motion correction to both the MR signal and the physiological slice timing vectors; however, this approach may introduce interpolation errors.

In conclusion, our simulations support that TRACC-PHYSIO is a practical and reliable time-domain approach to quantify cardiac and respiratory coupling, as well as their pulse delays with high millisecond-level temporal resolution. This method is particularly valuable in long-TR MRI acquisitions, where frequency-domain analyses are not feasible and physiological oscillations are obscured by cardiac aliasing. Its application to both existing and future datasets may yield important insights into the mechanisms of brain pulsations and their alterations in health and disease.

## Funding

Multiple National Institutes of Health grants supported this work, including funding from the National Institute of Aging: F30AG084336 (PI: Adam Wright) and RF1AG083762 (PI: Qiuting Wen). Support was provided by the National Institute of General Medical Sciences under Award Number T32GM148382.

## Data and code availability

All data and code are available upon request.

## Declaration of generative AI and AI-assisted technologies

During the preparation of this work, the authors used ChatGPT-4 for assistance in developing the code to simulate physiological signals (cardiac and respiratory) and minor components of the simulation design, including optimization of randomized signal noise generation and randomized physiological parameter selection. All code was thoroughly reviewed and edited by the authors, who take full responsibility for its content and implementation. ChatGPT-4 was also used to enhance the text’s language and readability. The authors reviewed and edited all text and take full responsibility for the content of the publication.

## Declaration of competing interests

The authors have no competing interests to disclose.

## Notes

### Competing Interest Statement

The authors have declared no competing interest.

## References

Bohr, T., Hjorth, P.G., Holst, S.C., Hrabětová, S., Kiviniemi, V., Lilius, T., Lundgaard, I., Mardal, K.A., Martens, E.A., Mori, Y., Nägerl, U.V., Nicholson, C., Tannenbaum, A., Thomas, J.H., Tithof, J., Benveniste, H., Iliff, J.J., Kelley, D.H., Nedergaard, M., 2022. The glymphatic system: Current understanding and modeling. iScience 25, 104987. 10.1016/J.ISCI.2022.104987

Huotari, N., Raitamaa, L., Helakari, H., Kananen, J., Raatikainen, V., Rasila, A., Tuovinen, T., Kantola, J., Borchardt, V., Kiviniemi, V.J., Korhonen, V.O., 2019. Sampling Rate Effects on Resting State fMRI Metrics. Front Neurosci 13. 10.3389/fnins.2019.00279

Iliff, J.J., Wang, M., Zeppenfeld, D.M., Venkataraman, A., Plog, B.A., Liao, Y., Deane, R., Nedergaard, M., 2013. Cerebral arterial pulsation drives paravascular CSF-Interstitial fluid exchange in the murine brain. Journal of Neuroscience 33. 10.1523/JNEUROSCI.1592-13.2013

Kiviniemi, V., Wang, X., Korhonen, V., Keinänen, T., Tuovinen, T., Autio, J., Levan, P., Keilholz, S., Zang, Y.F., Hennig, J., Nedergaard, M., 2016. Ultra-fast magnetic resonance encephalography of physiological brain activity-Glymphatic pulsation mechanisms? Journal of Cerebral Blood Flow and Metabolism 36, 1033–1045. 10.1177/0271678×15622047

Mestre, H., Tithof, J., Du, T., Song, W., Peng, W., Sweeney, A.M., Olveda, G., Thomas, J.H., Nedergaard, M., Kelley, D.H., 2018. Flow of cerebrospinal fluid is driven by arterial pulsations and is reduced in hypertension. Nat Commun 9. 10.1038/s41467-018-07318-3

Nair, V.V., Diorio, T.C., Wen, Q., Rayz, V.L., Tong, Y., 2024. Using respiratory challenges to modulate CSF movement across different physiological pathways: An fMRI study. Imaging Neuroscience 2, 1–14. 10.1162/imag_a_00192

Raitamaa, L., Huotari, N., Korhonen, V., Helakari, H., Koivula, A., Kananen, J., Kiviniemi, V., 2021. Spectral analysis of physiological brain pulsations affecting the BOLD signal. Hum Brain Mapp 42, 4298–4313. 10.1002/hbm.25547

Wen, Q., Muskat, J., Babbs, C.F., Wright, A.M., Zhao, Y., Zhou, X., Zhu, C., Tong, Y., Wu, Y.-C., Risacher, S.L., Saykin, A.J., 2025. Dynamic diffusion-weighted imaging of intracranial cardiac impulse propagation along arteries to arterioles in the aging brain. Journal of Cerebral Blood Flow & Metabolism. 10.1177/0271678×251320902

Wen, Q., Tong, Y., Zhou, X., Dzemidzic, M., Ho, C.Y., Wu, Y.-C., 2022. Assessing pulsatile waveforms of paravascular cerebrospinal fluid dynamics within the glymphatic pathways using dynamic diffusion-weighted imaging (dDWI). Neuroimage 260, 119464. 10.1016/j.neuroimage.2022.119464

Wen, Q., Wright, A., Tong, Y., Zhao, Y., Risacher, S.L., Saykin, A.J., Wu, Y., Limaye, K., Riley, K., 2023. Paravascular fluid dynamics reveal arterial stiffness assessed using dynamic diffusion-weighted imaging. NMR Biomed. 10.1002/nbm.5048

Wright, A., Xu, T., Koo, J., Zhao, Y., Tong, Y., Wen, Q., 2025. Age-related and sex-specific reduction in cerebral arterial-venous cardiac pulse delay in functional magnetic resonance imaging, in: International Society of Magnetic Resonance in Medicine. Hawaii.

Wright, A.M., Wu, Y.-C., Yang, H.-C., Risacher, S.L., Saykin, A.J., Tong, Y., Wen, Q., 2024. Coupled pulsatile vascular and paravascular fluid dynamics in the human brain. Fluids Barriers CNS 21, 71. 10.1186/s12987-024-00572-2

Zhang, J., Foster, E., Zhou, X., Wen, Q., 2025. Respiration is a driver for parenchyma hydrodynamics: insights from dynamic DWI, in: International Society of Magnetic Resonance in Medicine. Hawaii.

