## Supplemental for "TRACC-PHYSIO: Time-domain Resolution-Aligned Cross-Correlation to estimate PHYSIOlogical coupling and time delays in dynamic MRI"

**Supplemental Material:**

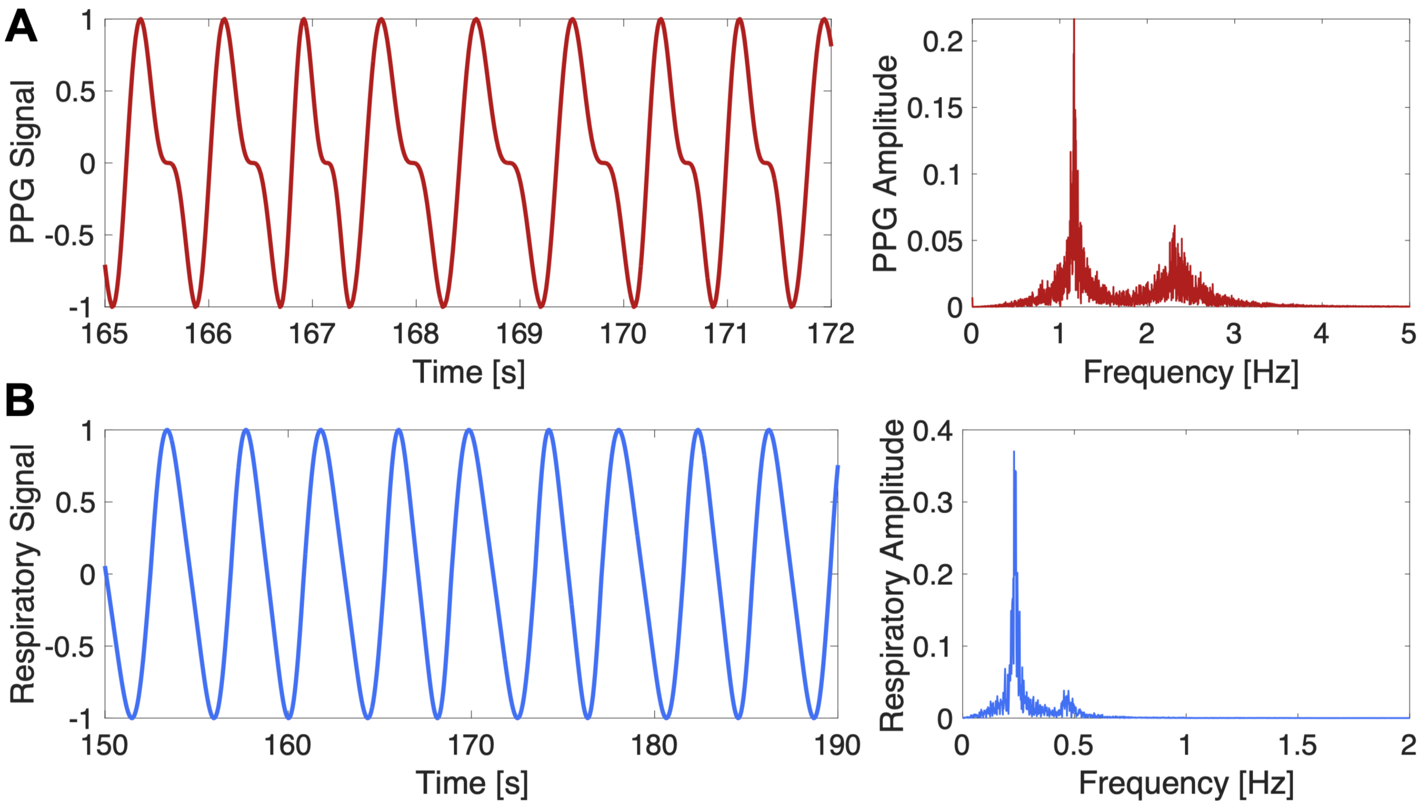


**Supplemental Figure 1:** Examples of the synthetic peripheral physiology waveforms. (**A**) Generated finger photoplethysmography (PPG) signal with a heart rate of 70 beats per minute and heart rate variability of 70 ms (left) along with its respective frequency spectrum (right). (**B**) Generated respiratory signal with a breathing rate of 14 breaths per minute and breathing rate variability of 300 ms (left) along with its respective frequency spectrum (right). **Note:** Synthetic physiological waveforms were generated with an initial length of 10 minutes, and the axes were zoomed to illustrate the waveform characteristics better. The physiological rate variability was defined as the standard deviation of the beat-to-beat (cardiac) or breath-to-breath (respiratory) intervals.

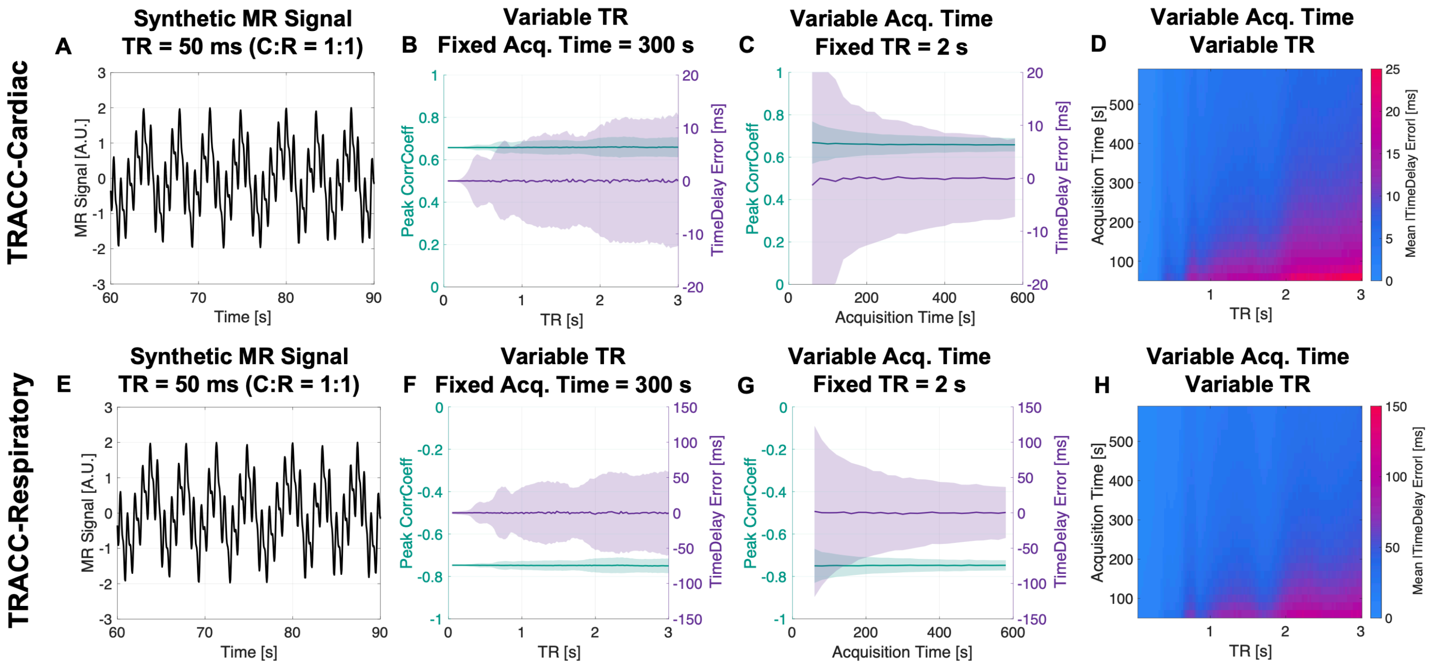


**Supplemental Figure 2: TRACC-PHYSIO demonstrated robust performance across varying TRs and acquisition times when the target had equal physiological component in the MR signal.** Top row (**A-D**): TRACC-Cardiac results for MR signals with C:R=1:1. Bottom row (**E-H**): TRACC-Respiratory results for MR signals with C:R=1:1. From left to right: (A & E) Example synthetic MR signals with TR = 50 ms. (**B & F**) The mean (solid line) and standard deviation (shaded) of the peak CorrCoeff and TimeDelay error with increasing TRs and a fixed acquisition time of 300 s. (**C & G**) The mean (solid line) and standard deviation (shaded) of the peak CorrCoeff and TimeDelay error with increasing acquisition time and a fixed TR of 2 s. (**D & H**) Heatmap of the mean absolute TimeDelay error across all combinations of TRs and acquisition times. **Note:** 5000 permutations were completed for each combination of TR and acquisition time. **Abbreviations:** C:R – cardiac-to-respiratory ratios.

To quantify CorrCoeff error, the ground truth was defined as the mean peak CorrCoeff at the shortest TR (0.05 s) and longest acquisition time (580 s). The mean peak CorrCoeff error was computed as the difference between the ground-truth and TRACC-estimated peak CorrCoeff. The mean absolute peak CorrCoeff error was minimal across all cardiac-respiratory ratios (C:R), TRs, and acquisition times in both TRACC-Cardiac (Supplemental Figure 3) and TRACC-Respiratory (Supplemental Figure 4).

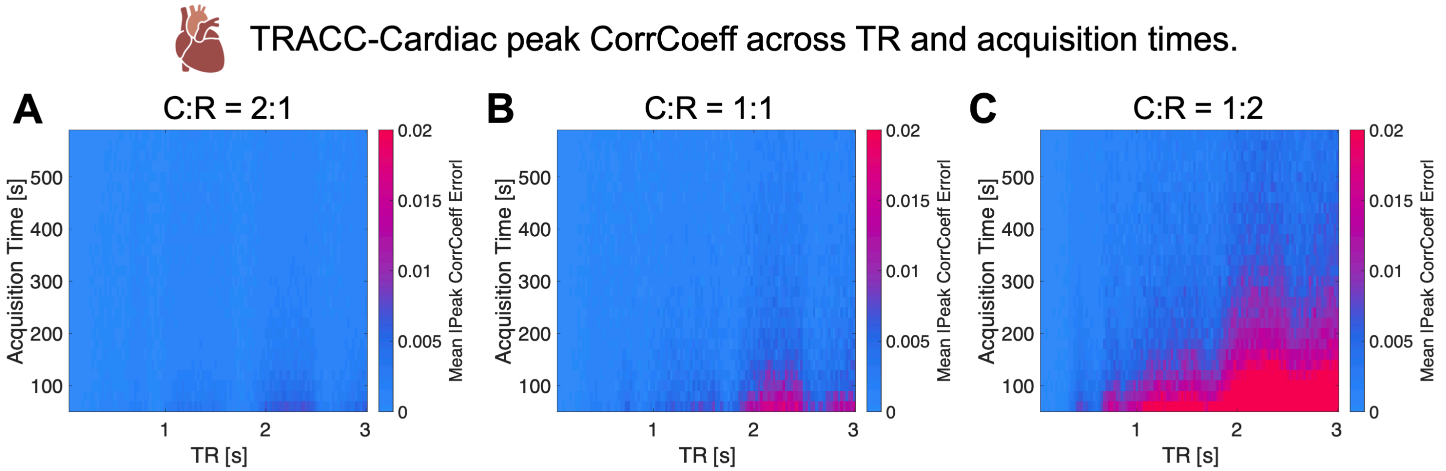


**Supplemental Figure 3:** Heatmaps of mean absolute peak CorrCoeff error in TRACC-Cardiac with varying MR repetition times (TR), acquisition times, and MR signal physiological components. The mean absolute peak CorrCoeff remained stable across large ranges of TRs and acquisition times in MR signals with different physiological components (**A**) C:R = 2:1, (**B**) C:R = 1:1, and (**C**) C:R = 1:2.

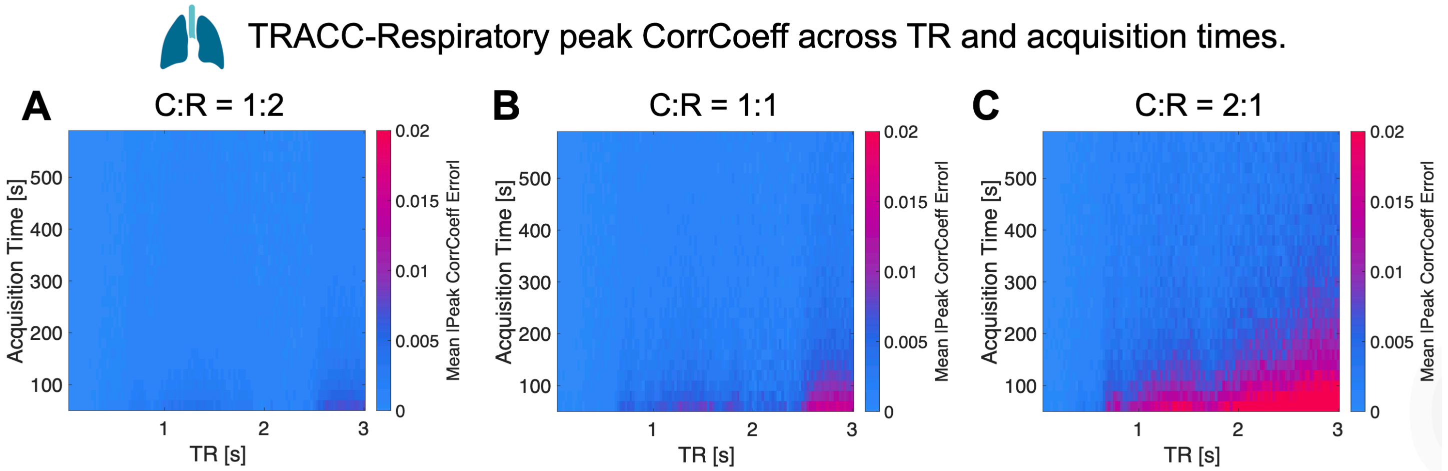


**Supplemental Figure 4:** Heatmap of the mean absolute peak CorrCoeff error in TRACC-Respiratory with varying MR repetition times (TR), acquisition times, and MR signal physiological components. The mean absolute peak CorrCoeff error remained stable across large ranges of TRs and acquisition times in MR signals with different physiological components (**A**) C:R = 1:2, (**B**) C:R = 1:1, and (**C**) C:R = 2:1.

### In-depth simulations of TRACC-PHYSIO with varying cardiac-to-respiratory ratios (C:R)

To assess how the cardiac-to-respiratory ratio (C:R) influence peak CorrCoeff and TimeDelay estimates, we performed simulations with varying C:R from 4:1 to 1:4 (in increments of 0.1; i.e. 4:1, 3.9:1, …, 1:1, 1:1.1, 1:1.2, …, 1:4) with a fixed acquisition time of 300 s and varying TR from 0.05 to 3 s (in increments of 0.05 s). For each combination of C:R and TR, 5,000 permutations were completed.

### TRACC-Cardiac Results:

Using TRACC-Cardiac, the peak CorrCoeff increased as the C:R increased (Supplemental Figure 5). With a fast TR of 50 ms, no TimeDelay errors were observed across all C:R (Supplemental Figure 5A). With a TR of 2 s, TimeDelay errors increased as the C:R decreased. At C:R < 1:3, the mean TimeDelay error deviated from 0, indicating a bias towards underestimation (Supplemental Figure 5B). Across all TRs, the estimation of the peak CorrCoeff was consistent for all C:R (Supplemental Figure 5C). As expected, TimeDelay errors increased as the cardiac component became less dominant (lower C:R ratio) and with longer TR.

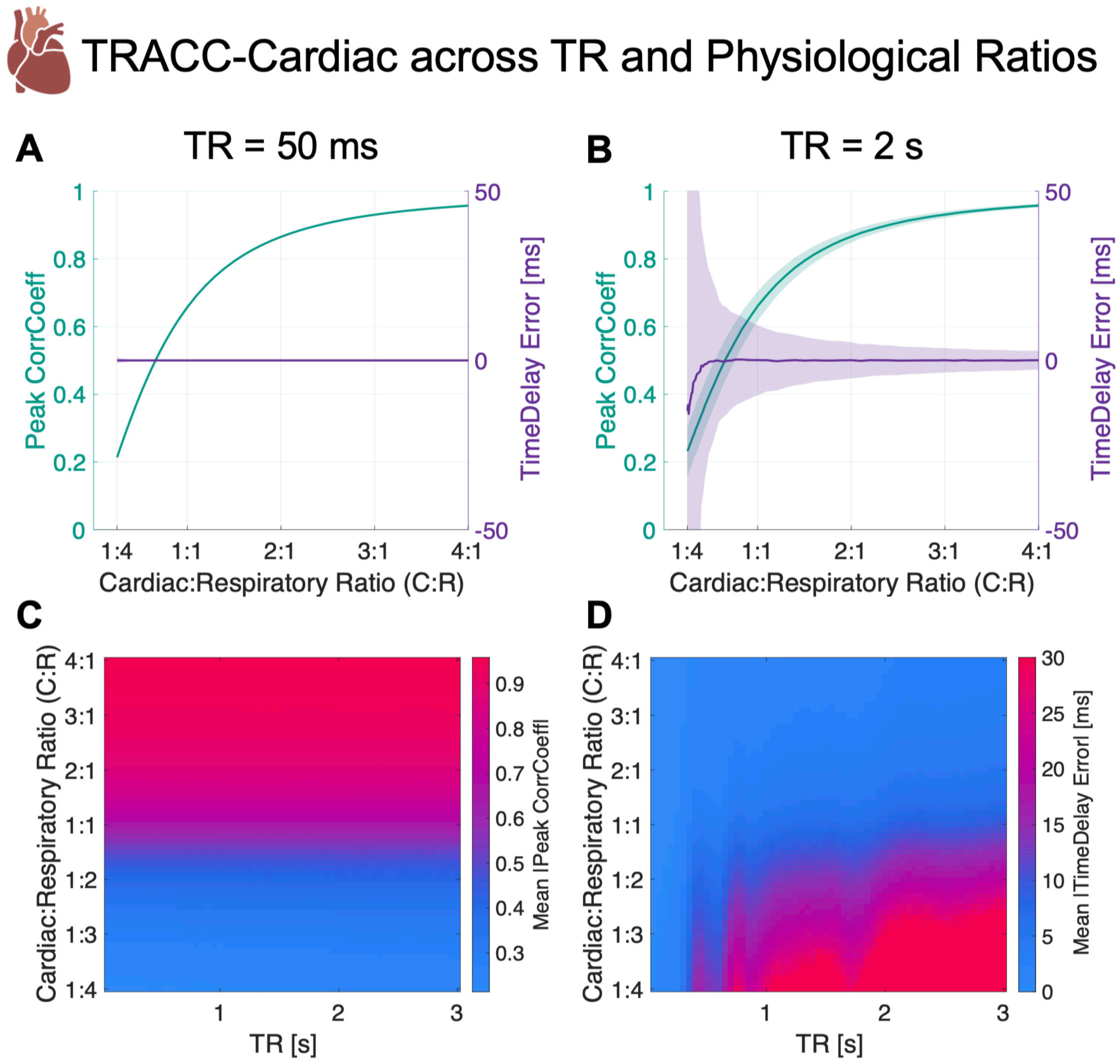


**Supplemental Figure 5:** The influence of the MR signal cardiac:respiratory ratio (C:R) and repetition time (TR) in TRACC-Cardiac measurements, with a fixed acquisition time of 300 s. The peak CorrCoeff and TimeDelay error across varying C:R at, (**A**) TR = 50 ms and (**B**) TR = 2 s. (**C**) The mean peak CorrCoeff and (**D**) mean TimeDelay error across each combination of C:R and TR.

Variation in MR signal physiological components introduced changes in the peak CorrCoeff, which initially increased nonlinearly and then gradually plateaued as the C:R ratio increased. Since the peak CorrCoeff reflects the dominance of the physiological signal—and this dominance influences TimeDelay accuracy—CorrCoeff may serve as an indicator of confidence in the TimeDelay estimate. To evaluate the TimeDelay error vs peak CorrCoeff relationship, we fit a linear mixed-effects model to assess the relationship between the absolute TimeDelay error, the absolute peak CorrCoeff, and TR. Each of the 5000 permutations for each (TR, Physiological Ratio) pair were treated as a repeated measure (Repeat), using Equation 1.

$$|\text{TimeDelay Error}| \sim 1 + |\text{Peak CorrCoeff}| \times \text{TR} + (1 \mid \text{Repeat}) \quad (1)$$

The regression results showed that absolute TimeDelay error was negatively associated with absolute peak CorrCoeff and positively associated with TR (Supplemental Table 1). A significant negative interaction between peak CorrCoeff and TR indicated that the reduction in TimeDelay error observed with high peak CorrCoeff was stronger at longer TRs. In other words, at longer TRs, the peak CorrCoeff was a stronger predictor of TimeDelay error.

**Supplemental Table 1:** The relationship of absolute TimeDelay error with the absolute peak CorrCoeff and repetition time (TR) determined with a linear mixed effects model using every simulated combination of physiological ratio and TR with TRACC-Cardiac.

| Term | Estimate (95% CI) | SE | t-stat | p-value |
|---|---|---|---|---|
| (Intercept) | 11.7 (10.6, 12.9) | 0.59 | 19.8 | p<0.001 |
| \|Peak CorrCoeff\| | -15.9 (-16.8, -15.0) | 0.45 | -35.2 | p<0.001 |
| TR | 33.8 (33.2, 34.5) | 0.33 | 103.6 | p<0.001 |
| \|Peak CorrCoeff\| × TR | -44.6 (-45.0, -44.1) | 0.21 | -209.7 | p<0.001 |

**TRACC-Respiratory Results:**

Using TRACC-Respiratory, the peak CorrCoeff increased as C:R decreased (i.e., the respiratory component increased, Supplemental Figure 5). With a fast TR of 50 ms, no TimeDelay errors were experienced across all C:R (Supplemental Figure 5A). With a TR of 2 s, variations in TimeDelay errors increased as C:R increased, and no significant error bias was observed at any C:R (Supplemental Figure 5B). Across all TRs, the estimation of the peak CorrCoeff was consistent for all C:R (Supplemental Figure 5C). Minimal TimeDelay errors were observed with TRs<600 ms for all C:R (Supplemental Figure 5D). For TRs>600 ms, minimal TimeDelay errors were observed in signals with equal physiological components (C:R = 1) or signals dominated by respiratory amplitude (C:R<1, Supplemental Figure 5D). Increased TimeDelay errors were present when TRs exceeded 1 s and C:R>2 (Supplemental Figure 5D). As expected, TimeDelay errors increased as the respiratory component was less dominant (higher C:R ratio) and with longer TR.

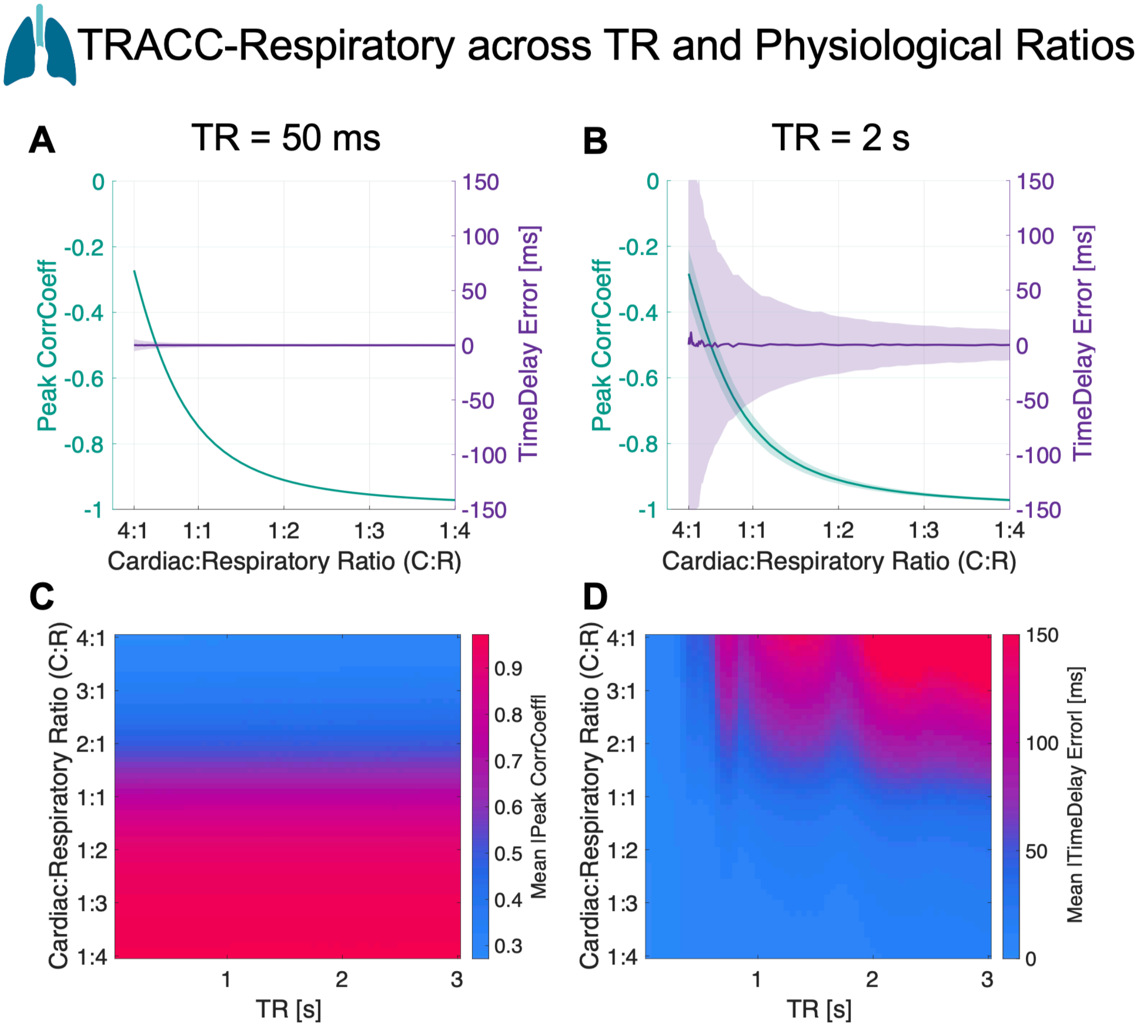


**Supplemental Figure 6:** The influence of the MR signal cardiac:respiratory ratio (C:R) and repetition time (TR) in TRACC-Respiratory measurements, with a fixed acquisition time of 300 s. The peak CorrCoeff and TimeDelay error across varying C:R at, (**A**) TR = 50 ms and (**B**) TR = 2 s. (**C**) The mean peak CorrCoeff and (**D**) mean TimeDelay error across each combination of C:R and TR.

The modeling of TimeDelay errors in TRACC-Respiratory behaved similarly to TRACC-Cardiac. The absolute TimeDelay error was negatively associated with absolute peak CorrCoeff and positively associated with TR. Additionally, a negative interaction between peak CorrCoeff and TR indicated that the reduction in TimeDelay error with higher peak CorrCoeff was more pronounced at longer TRs (Supplemental Table 2). In other words, at longer TRs, the peak CorrCoeff was a stronger predictor of TimeDelay error.

**Supplemental Table 2:** The relationship of absolute TimeDelay error with the absolute peak CorrCoeff and repetition time (TR) determined with a linear mixed effects model using every simulated combination of physiological ratio and TR with TRACC-Respiratory.

| **Term** | **Estimate (95% CI)** | **SE** | **t-stat** | **p-value** |
|---|---|---|---|---|
| (Intercept) | 38.6 (37.0, 40.2) | 0.80 | 48.2 | p<0.001 |
| \|Peak CorrCoeff\| | -40.5 (-42.3, -38.6) | 0.96 | -42.0 | p<0.001 |
| TR | 72.5 (71.7, 73.3) | 0.41 | 177.4 | p<0.001 |
| \|Peak CorrCoeff\| × TR | -73.0 (-73.9, -72.1) | 0.46 | -158.5 | p<0.001 |
